# Neuroblastoma-derived small extracellular vesicles retain cellular identity and adrenergic/mesenchymal state signatures

**DOI:** 10.64898/2026.09.21.753086

**Authors:** Maddalena Grimaldi, Marieluise Kirchner, Annabell Szymansky, An Hendrix, Agnieszka Münster-Wandowski, Holger Gerhardt, Anja H. Hagemann, Marco Lodrini, Martin Auber, Philipp Mertins, Hedwig E. Deubzer

**Affiliations:** Department of Pediatric Oncology and Hematology, Charité – Universitätsmedizin Berlin, Berlin, Germany; Proteomics Platform, Max-Delbrück-Center for Molecular Medicine (MDC) in the Helmholtz Association, Berlin, Germany; Berlin Institute of Health (BIH) at Charité, Berlin, Germany; German Cancer Consortium (DKTK), Partner Site Berlin, and German Cancer Research Center (DKFZ), Heidelberg, Germany; Department of Human Structure and Repair, Laboratory of Experimental Cancer Research, Ghent University, Ghent, Belgium; Cancer Research Institute Ghent (CRIG), Ghent University, Ghent, Belgium; Institute of Integrative Neuroanatomy, Charité - Universitätsmedizin Berlin, Berlin, Germany; Max-Delbrück-Center for Molecular Medicine (MDC) in the Helmholtz Association, Berlin 13125, Germany; German Center for Cardiovascular Research (DZHK) Partner site Berlin, Berlin 10785, Germany

**Keywords:** Extracellular vesicles, mass spectrometry, proteomics, adrenergic-mesenchymal plasticity, premetastatic niche formation, intercellular communication, microenvironment

## Abstract

Neuroblastoma is a heterogeneous pediatric malignancy characterized by distinct genomic subgroups and cell-state plasticity between adrenergic (ADRN) and mesenchymal (MES) phenotypes. While small extracellular vesicles (sEVs) mediate pre-metastatic niche priming, the degree to which sEV proteomes reflect parental genomic and phenotypic identity remains unresolved. Here, we isolated sEVs from six neuroblastoma cell models (IMR-5/75, BE(2)-C, GI-ME-N, CLB-GA, LAN-6, and SK-N-FI) representing three genomic subgroups (MYCN-amplified, TERT-rearranged, and unaltered) using OptiPrep density gradient centrifugation followed by size exclusion chromatography and ultrafiltration. High-resolution mass spectrometry of sEV proteomes revealed that sEVs carry highly reproducible protein signatures that tightly correlate with the parental genomic subgroups. Analyzing paired cell/sEV samples, we demonstrate cell-to-EV concordance of lineage-specific markers: ADRN sEVs selectively enriched L1CAM and dopamine β-hydroxylase (DBH), whereas MES sEVs enriched SERPINE1 and type III collagen (COL3A1). These findings indicate that neuroblastoma-derived sEVs systematically preserve both the genomic landscape and the lineage transdifferentiation states of their cells of origin. This study establishes a robust molecular foundation for utilizing sEV-based proteomics as a surrogate to track cellular plasticity and for studying sEV-mediated microenvironmental remodeling.

## INTRODUCTION

Neuroblastoma is the most common extracranial solid tumor of childhood, arising from sympathoadrenal progenitor cells of the neural crest.^1^ It exhibits a remarkably heterogeneous clinical spectrum, ranging from spontaneous regression in infants to highly aggressive, refractory progression in older children.^2^ High-risk neuroblastoma represents approximately 50% of cases and is characterized by a dismal 5-year survival rate of only ∼50%, primarily due to emerging therapeutic resistance and metastatic burden.^3^ Neuroblastoma exhibits an organotropism, seeding into locoregional lymph nodes, infiltrating the bone marrow and spreading into liver, lung and, more seldomly, the central nervous system.

Before circulating neuroblastoma cells can successfully colonize distant, specialized tissues, the primary tumor is thought to orchestrate the formation of a supportive, tumor-promoting microenvironment known as the pre-metastatic niche.^4,5^ Recent advances have highlighted small extracellular vesicles (sEVs) as systemic mediators of this long-range intercellular communication.^4,6^ Released in large quantity, sEVs actively home to future metastatic organs.^4,5^ At these metastatic sites, sEV-associated cargoes initiate a cascade of pathological remodeling steps, including the induction of vascular permeability, extracellular matrix remodeling - exemplified by fibronectin deposition - and stromal activation.^4,5^ Furthermore, neuroblastoma-derived sEVs play a role in systemic immune evasion and local immunosuppression within the niche.^4,7^ They carry potent immunomodulatory molecules such as PD-L1 and HLA-G that inhibit cytotoxic T-cell proliferation and can actively reprogram local bone marrow macrophages toward tumor-supportive, M2-like phenotypes.^4,7^

Biological complexity and treatment resistance of high-risk neuroblastoma are compounded by cellular and lineage plasticity.^8-13^ Transcriptomic and epigenetic profiling has established that neuroblastoma cells exist in two distinct, interconvertible differentiation states: a highly differentiated adrenergic (ADRN) phenotype and an immature, neural crest-like mesenchymal (MES) phenotype.^14-16^ While ADRN cells dominate primary tumor masses, the MES phenotype exhibits enhanced migratory capacity, stemness properties, and profound resistance to standard chemotherapeutic regimens. Recent studies indicate that this cellular state is also reflected in the molecular envelopes of secreted sEVs as distinct microRNA and protein signatures have been identified in sEVs derived from ADRN and MES subtypes.^17,18^ This may provide a liquid-biopsy surrogate to monitor the dynamic phenotypic evolution of the tumor.

However, despite our understanding of the general role of sEVs in mediating premetastatic niche formation and reflecting lineage identity, a comprehensive, high-resolution comparative map of the proteomic cargo of paired neuroblastoma cells and their corresponding sEVs remains missing. Specifically, it remains unclear to what extent the proteomic envelope of secreted sEVs reflects the precise genomic landscape and ADRN vs. MES phenotypes of their donor cells. To address this gap, we systematically profiled the proteomic cargo of sEVs and paired cells across a panel of neuroblastoma cell models utilizing high-throughput mass spectrometry. We show that the proteomic cargo of secreted sEVs correlates with the genomic profile of the parental cell. Notably, we also demonstrate that the mesenchymal-to-adrenergic trajectory is preserved and readable within the proteomic cargo of secreted sEVs.

## MATERIALS AND METHODS

### Cell culture

The BE(2)-C cell line (RRID: CVCL_0529) was obtained from ECACC (Salisbury, UK) and the GI-ME-N cell line (RRID: CVCL_1232) from the DSMZ (Braunschweig, Germany). CLB-GA (RRID: CVCL_9529), IMR-5/75 (RRID: CVCL_M473) and LAN-6 (RRID: CVCL_1363) cell lines were kindly provided by M. Fischer (University Hospital Cologne), the SH-EP (RRID:CVCL_0524; epithelial-like subclone of SK-N-SH) and SK-N-AS (RRID: CVCL_1700) cell lines by L. Savelyeva (German Cancer Research Center (DKFZ), Heidelberg) and the SK-N-FI cell line (RRID: CVCL_1702) by J. Schulte (Charité, Berlin). Cell lines were routinely maintained at 37 °C and 5% CO^2^ in Gibco DMEM (Thermo Fisher Scientific, Dreieich, Germany) supplemented with 10% fetal calf serum (Merck) and 1% non-essential amino acids (Lonza, Cologne, Germany). Cell lines were authenticated using high-throughput SNP-based assays^19^ and regularly monitored for mycoplasma using PlasmoTest (InvivoGen, San Diego, CA, U.S.A.) according to the manufacturer’s instructions.

### Preparation of conditioned medium for small extracellular vesicle isolation

EV-depleted fetal calf serum (Thermo Fisher Scientific) was used as a protein source for experiments intended to obtain conditioned medium suitable for extracellular vesicle purification. In brief, cell cultures were washed three times using DMEM followed by a 24 h incubation with EV-harvesting medium (DMEM supplemented with 10% EV-depleted fetal calf serum and 1% non-essential amino acids). Conditioned medium was harvested, centrifuged at 200 x *g* for 10 min at 4 °C and passed through an 0.45 *μ*m cellulose acetate filter (Corning, Corning, NY, U.S.A.) to remove cell debris.^20^ The conditioned medium was concentrated approximately 200-fold at 4 °C using a 10 kDa Centricon Plus-70 centrifugal filter device (Merck Millipore, Burlington, MA, U.S.A.).^20^ Following collection of conditioned medium, adherent neuroblastoma cells were trypsinized, and cell viability was assessed using a VI-CELL-XR Cell Viability Analyzer (Beckman Coulter, Brea, CA, U.S.A.) based on the trypan blue-exclusion method. Cultures with cell viability >85% were included in further analyses.

### OptiPrep density gradient centrifugation

A discontinuous iodixanol gradient was used for sEV isolation as described^21^ with minor modifications. In brief, an iodixanol working solution was prepared by combining a stock solution of OptiPrep™ (60% (w/v) aqueous iodixanol solution; Sigma-Aldrich) with solution buffer (0.25 M sucrose, 6 mM EDTA, 60 mM Tris-HCl, pH 7.4). This working solution was mixed with appropriate amounts of homogenization buffer (0.25 M sucrose, 1 mM EDTA, 10 mM Tris-HCl, pH 7.4) to obtain 5%, 10%, 20% and 40% iodixanol solutions. The gradient was formed by sequentially layering 3 mL of 40%, 3 mL of 20%, 3 mL of 10% and 2.5 mL of 5% solutions on top of each other in a 14 mL Ultra-Clear™ open top centrifugal tube (Beckman Coulter). A total of 700 *µ*L concentrated conditioned cell culture medium was overlaid on the top of the gradient followed by ultracentrifugation at 100,000 x *g* for 18 h and 4 °C using a SW 40 Ti rotor (Beckman Coulter). Gradient fractions of 1 mL were collected from the top of the gradient, and EV-enriched fractions 5, 6 and 7 were pooled and further processed through size-exclusion chromatography.

### Size exclusion chromatography and ultrafiltration

Sepharose CL-2B (GE Healthcare, Chicago, IL, U.S.A.) was washed three times with PBS containing 0.32% (w/v) trisodiumcitrate dehydrate (Carl Roth, Karlsruhe, Germany).^22^ To obtain columns for size exclusion chromatography, 10 mL sepharose CL-2B was stacked into a 10 ml syringe (Neolab, Heidelberg, Germany) containing a nylon net with 20 *µ*M pore size (Merck Millipore) at the bottom.^22^ Altogether, 2 mL of sample were loaded, and 1 mL fractions of eluate were collected and pooled. Eluates were centrifuged in a swinging bucket rotor equipped with an Amicon Ultra-2 10 k centrifugal filter (Merck Millipore) at 3,000 x g and 4 °C for 10 min to obtain a final volume of 100 -200 *µ*l purified EV eluate in PBS. Samples were stored at -80 °C.

### Nanoparticle tracking analysis

Nanoparticle tracking analysis was performed using a NanoSight LM-10 microscope equipped with a 488 nm laser (Malvern Panalytical, Malvern, UK). Samples were diluted with filtered PBS to a particle concentration ranging between 3×10^8^ and 1×10^9^ particles/mL. In total, three 60-second videos were recorded for each sample with camera level 14 and detection threshold 3.^22^ Recorded videos were analyzed with NTA Software version 3.2 (Malvern Panalytical).

### Whole-mount immunoelectron microscopy

The sEV suspension was fixed in 2% paraformaldehyde and deposited on Formvar carbon-coated glow-discharged nickel grids (Plano, Wetzlar, Germany) for investigation by whole-mount immunoelectron microscopy as described.^23^ Grids were examined in a Zeiss transmission electron microscope equipped with a ProScan 2K slow-scan CCD camera controlled by the ImageSP software (Tröndle, Moorenweis, Germany). For negative controls, primary antibodies were omitted.

### Western blotting

sEV suspensions and cells were lysed for western blotting in buffer containing 20 mM Tris-HCl, 7 M urea, 0.01 % Triton X-100, 40 mM MgCl2, Complete^®^ protease inhibitor cocktail (Roche, Mannheim, Germany) and 100 mM DTT as reducing agent if indicated. The following antibodies were used: mouse monoclonal anti-ALIX (1:1000; Cell Signaling, Danvers, MA, U.S.A.), rabbit polyclonal anti-CALR (1:1000; Cell Signaling), anti-CD63 (1:250; Thermo Fisher Scientific), mouse monoclonal anti-CD81 (1:300; Thermo Fisher Scientific), mouse monoclonal anti-COL3A1 (1:500; Santa-Cruz Biotechnology, Dallas, TX, U.S.A.), mouse monoclonal anti-DBH (1:800; Santa-Cruz Biotechnology), mouse monoclonal anti-FLOT1 (1:1000; BD Biosciences, San Jose, CA, U.S.A), mouse monoclonal anti-GAPDH (1:20000; Merck Millipore), mouse monoclonal mouse monoclonal anti-L1CAM (1:1000; Thermo Fisher Scientific) and rabbit polyclonal anti-SERPINE1 (1:1000, Novus Biologicals, Littleton, CO, U.S.A.).

### Proteome analyses of sEVs and cells by mass spectrometry

sEV suspensions were mixed 1:1 with SDC lysis buffer (4% sodium dexoxycholate, 200 mM Tris pH 8.0, 2 mM EDTA, 200 mM NaCl, 20 mM DTT, 80 mM chloroacetamide) and heated at 95 °C for 10 min. Samples were treated with 25 U benzonase (Merck Millipore) for 30 min at room temperature followed by quenching with 40 mM DTT. Single-pot, solid-phase-enhanced sample-preparation (SP3)-based protein enrichment, clean-up and tryptic digest were performed as described.^24^ Briefly, samples were incubated for 20 min with 100 *µ*g SP3 paramagnetic beads (Promega, Madison, WI, U.S.A.) and acetonitrile (70% final concentration) followed by two washing steps with 70% EtOH and one washing step with 100% acetonitrile using a magnetic rack. Beads were air-dried, then 200 ng sequence-grade trypsin (Promega, Madison, WI, U.S.A.) and 200 ng lysyl endopeptidase LysC (Wako Chemicals, Neuss, Germany) dissolved in 30 *µ*l of 50 mM ammonium bicarbonate were added. Digest was performed on a thermo shaker overnight at 37 °C. Peptide containing supernatants were collected, and beads were incubated twice with 50 *µ*l of 50 mM ammonium bicarbonate. Corresponding supernatants were combined, and formic acid was added to obtain a final concentration of 1%. Peptides were stored on StageTips.^25^

Cell pellets were resuspended in SDC lysis buffer (1% sodium deoxycholate, 100 mM Tris-HCl pH 8.0, 1 mM EDTA, 150 mM NaCl, 10 mM DTT, 40 mM chloroacetamide), heated at 95°C for 10 min, cooled down to room temperature and incubated with 100 U benzonase (Merck Millipore) for 30 min. After centrifugation at 18,000 g for 15 min at 4 °C, supernatants were collected and subjected to protein quantification using the Bio-Rad DC Protein assay (Bio-Rad, Hercules, CA, U.S.A.). In total, 100 *µ*g protein were digested in-solution using 2 *µ*g sequence-grade trypsin and 2 *µ*g lysyl endopeptidase LysC overnight at 37 °C. The reaction was stopped by adding trifluoroacetic acid (final concentration 1%), and peptides were desalted and cleaned-up using the StageTips protocol.^25^

Peptides were eluted from the stage tips using 80% acetonitrile and 0.1% formic acid. After evaporation, the organic solvent peptides were resolved in sample buffer (3% acetonitrile, 0.1% formic acid). An analytical run with injection of 1 *µ*g peptide was performed for each sample. Peptides were separated on a 20 cm reversed-phase column (75 *µ*M inner diameter packed with ReproSil-Pur C18-AQ, particle size 1.9 *µ*M; Dr. Maisch, Ammerbuch-Entringen, Germany) of a high-performance liquid chromatography system (Thermo Fisher Scientific). The gradient was run for 200 min with a flow rate of 250 nL/min and buffer B concentrations increasing from 2% to 60%. Peptides were measured on a Q Exactive HF-X instrument (Thermo Fisher Scientific). The mass spectrometer was operated in the data dependent mode with a 70K resolution, 3 x 10^6^ ion count target and maximum injection time of 10 ms for the full scan, followed by Top 20 MS2 scans with 15K resolution, 1 x 10^5^ ion count target and maximum injection time of 22 ms.

Raw data were processed using the MaxQuant software package v1.6.3.4.^26^ The Andromeda search machine^27^ included in MaxQuant v.1.6.3.4 was employed to map the MS2 spectra against the HUMAN.2019-17 UniProt database including isoform annotations and containing both forward and reverse sequences. The search included variable modifications of oxidation (M), N-terminal acetylation, deamidation (N and Q) and fixed modifications of carbamidomethyl cysteine. Minimal peptide length was defined as seven amino acids, and a maximum of three missed cleavages was allowed. The false discovery rate was set to 1% both for peptide and protein identification. Unique and razor peptides were considered for quantification. Retention times were recalibrated based on the built-in nonlinear time-rescaling algorithm. MS2 identifications were transferred between runs for sEV and cell samples separately using the “match between runs” algorithm. The integrated LFQ quantification algorithm^28^ was likewise applied separately for sEV and cell samples. The resulting text files were filtered to exclude reverse database hits, potential contaminants and proteins only identified by site.

### Statistical analysis

sEV and cell proteome data were analyzed using the Perseus software v.1.6.2.1^26^ and R studio version 1.1.463 with R versions 3.5.1^29^ and 3.6.1^30^. Label-free quantification intensity values were filtered for a minimum value of four in at least one group. After log2 transformation, missing values were imputed with random noise simulating the detection limit of the mass spectrometer (log-normal distribution with 0.25 x SD of logarithmized values, downshifted by 1.8 x SD).^31,32^ Statistical data analysis was performed using Welch’s t-test to analyze differences in the proteome cargo of sEVs for each cell line as well as among genomic subgroups. The Benjamini-Hochberg procedure^33^ was applied for multiple testing correction, with significance cut-offs set to FDR < 1% for genomic and cell line comparisons, or FDR < 5% for cell-line-specific enrichment of proteins in parental cells and sEVs. Heatmaps were created using the heatmap package, version 1.0.13, in R.^34^

To ensure that biological conclusions were not technical artifacts of the mass spectrometry missing value imputation algorithm, we conducted a sensitivity analysis on the primary lineage markers using the non-imputed data matrix. We verified that the key lineage-state indicators (L1CAM, DBH, SERPINE1, and COL3A1) were robustly and abundantly detected (spectral counts and raw LFQ intensities) in all biological replicates of their respective positive cell states, while being significantly lower abundant or absent (below the analytical detection limit) in the opposite states (**Suppl. Fig. S1**). Thus, their statistical identification as highly differentially abundant cargo is mathematically independent of the imputation parameters employed in Perseus. Heatmaps were created using the pheatmap package, version 1.0.13. in R.^34^

### EV-TRACK

All relevant experimental data have been submitted to the EV-TRACK knowledgebase^35^ (EV-TRACK ID will be provided).

## RESULTS

### Isolation and characterization of extracellular vesicles from neuroblastoma cell lines

To establish a panel of neuroblastoma-derived extracellular vesicles for downstream analyses, sEVs were isolated by OptiPrep density gradient centrifugation followed by size-exclusion chromatography and ultrafiltration from conditioned medium of six neuroblastoma cell lines: IMR-5/75, BE(2)-C, GI-ME-N, CLB-GA, LAN-6 and SK-N-FI (**Fig. 1**). Particle-size analysis showed heterogeneous populations of predominantly small particles in preparations from all six cell lines (**Fig. 1A**). Mean particle diameters ranged from 105.4 ± 2.2 nm in BE(2)- C to 160.7 ± 3.1 nm in LAN-6, whereas modal diameters ranged from 74.1 ± 3.2 nm in CLB-GA to 114.0 ± 5.5 nm in SK-N-FI (mean ± SEM, *n* = 5 independent sEV isolations). The mean diameters for IMR-5/75, BE(2)-C, GI-ME-N, CLB-GA, LAN-6 and SK-N-FI were 145.7 ± 4.1, 105.4 ± 2.2, 144.7 ± 4.7, 130.3 ± 4.1, 160.7 ± 3.1 and 156.1 ± 2.3 nm, respectively, with corresponding modal diameters of 94.5 ± 5.9, 76.3 ± 4.2, 102.8 ± 3.2, 74.1 ± 3.2, 102.4 ± 5.6 and 114.0 ± 5.5 nm. Transmission electron microscopy further showed nanoscale vesicular structures with heterogeneous morphology and size, consistent with the particle distributions measured by NTA (**Fig. 1B**). To assess the biochemical distribution of sEV-associated proteins across the SEC eluate, fractions 4-9 were analyzed individually by Western blot analysis (**Fig. 1C**). ALIX, FLOT1, CD63 and CD81 were detected across the early SEC fractions, whereas CALR was not detected in any of fractions 4-9. In contrast, calreticulin was readily detectable in the corresponding whole-cell lysates, supporting the separation of sEV-enriched material from cellular components. On the basis of the particle characteristics and sEV-marker distribution, fractions 5, 6 and 7 were selected and pooled for all subsequent experiments. Rank abundance distribution and marker intensity analysis employing the proteomic data verified the quality of the isolated sEVs, characterized by high abundance of positive sEV markers and low or undetectable expression of cellular contamination markers (**Suppl. Fig. S2**). Together, these data established SEC fractions 5-7 as sEV-enriched preparations from each of the six neuroblastoma cell lines.

**Fig 1.**
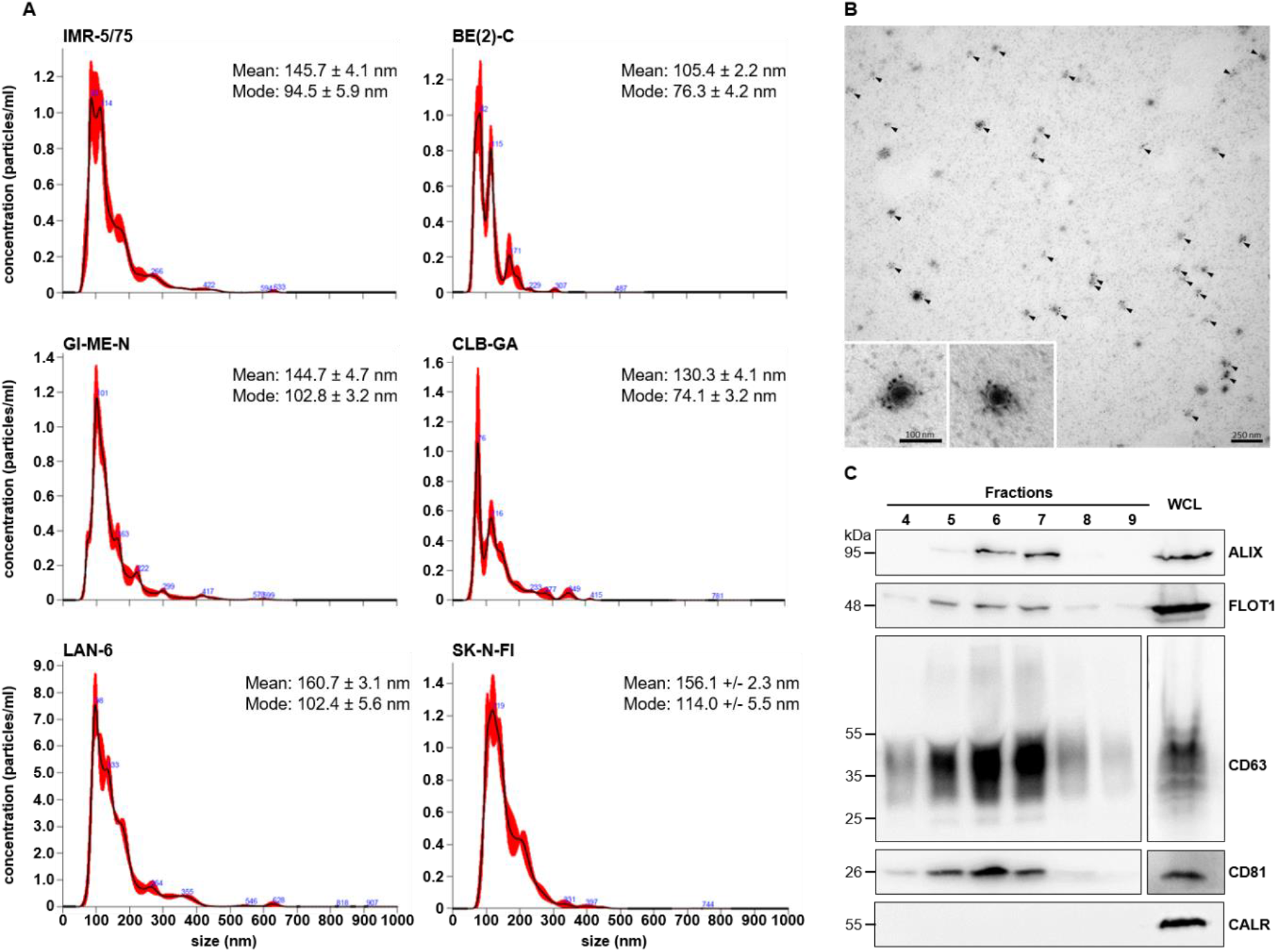
Isolation and characterization of extracellular vesicles from neuroblastoma cell lines. **A**, Particle-size distributions of sEVs isolated by size-exclusion chromatography (from conditioned medium of IMR-5/75, BE(2)-C, GI-ME-N, CLB-GA, LAN-6 and SK-N-FI neuroblastoma cells, determined by the NanoSight LM-10 microscope. Mean and modal particle diameters are shown as mean ± SEM from five independent sEV isolations (*n* = 5). **B**, Representative transmission electron microscopy image of SEC-isolated EVs. Arrowheads indicate vesicular structures. Insets show representative particles at higher magnification. Scale bars, 250 nm and 100 nm, as indicated. **C**, Western blot analysis of SEC fractions 4-9 for ALG-2-interacting protein X (ALIX), flotillin (FLOT1), CD63, CD81 and calnexin (CAL). Corresponding whole-cell lysates (WCL) are shown in the separate panel to the right. Fractions 5-7 were pooled for subsequent experiments.

### Neuroblastoma genomic subgroups are reflected in the sEV proteome

We next investigated whether the proteomic composition of neuroblastoma-derived sEVs reflected the genomic characteristics of the respective parental cells. The six neuroblastoma cell lines represented three genomic groups^36^: *MYCN*-amplified IMR-5/75 and BE(2)-C cells, *TERT*-rearranged GI-ME-N and CLB-GA cells, and LAN-6 and SK-N-FI cells, which lacked both *MYCN* amplification and *TERT* rearrangement (**Fig. 2**). After data filtering (see Material and Method section), a total of 3194 proteins were used for comparison of each cell line to all others, identifying 1409 differentially expressed proteins (DEPs) in total, and revealing distinct sEV protein profiles across the six neuroblastoma cell lines. (**Fig. 2**). Noteworthy, GI-ME-N shows strongest enrichment signal (log2 FC >0) and IMR-5/75 strongest depletion signal (log2 FC < 0) in the sEV proteome. To determine whether these profiles contained features shared by cell lines with the same genomic characteristics, we compared differentially abundant proteins between the genomic groups, resulting in 144 significantly different proteins in total. IMR-5/75 and BE(2)-C shared a set of differentially abundant sEV proteins (110 DEPs), consistent with a common protein signature among the *MYCN*-amplified cell lines. In contrast to the *MYCN*-amplified subgroup, the TERT-rearranged lines (GI-ME-N and CLB-GA) displayed a remarkably small overlapping set of differentially abundant sEV proteins, sharing only 11 DEPs in the subgroup comparison. This striking discordance presents a critical biological finding. The proteomic profiles of GI-ME-N and CLB-GA sEVs diverged profoundly despite sharing the same TERT-rearranged genomic driver. We hypothesized that this discordance was driven by an underlying lineage-state confounder, as CLB-GA cells display a highly differentiated adrenergic (ADRN) phenotype, whereas GI-ME-N represents a mesenchymal (MES) phenotype within the discovery panel. To isolate true TERT-associated features from this cell-state background, we performed a supplementary sub-analysis. We compared the proteome of CLB-GA sEVs (ADRN) exclusively with the other ADRN sEV preparations (IMR-5/75, BE(2)-C, LAN-6, SK-N-FI), thereby filtering out cell-state-associated variance. This targeted comparison successfully isolated 43 sEV proteins that were specifically enriched in CLB-GA, which represent candidate markers associated with TERT-rearrangement in an adrenergic background. Conversely, the unique proteomic features of GI-ME-N sEVs (which displayed the strongest enrichment signal with log2 FC > 0 in the overall panel) were heavily dominated by mesenchymal differentiation signatures rather than TERT status. This analysis demonstrates that lineage identity and cell-state differentiation exert a more dominant influence on the sEV proteome than specific oncogenic genomic alterations, necessitating cell-state stratification in downstream diagnostic development. LAN-6 and SK-N-FI, which lacked both genomic alterations, also shared a distinct set of differentially abundant proteins (26 DEPs). Thus, in addition to cell line-specific differences, sEV protein composition exhibited common features among neuroblastoma cells sharing the same major genomic characteristics.

**Fig 2.**
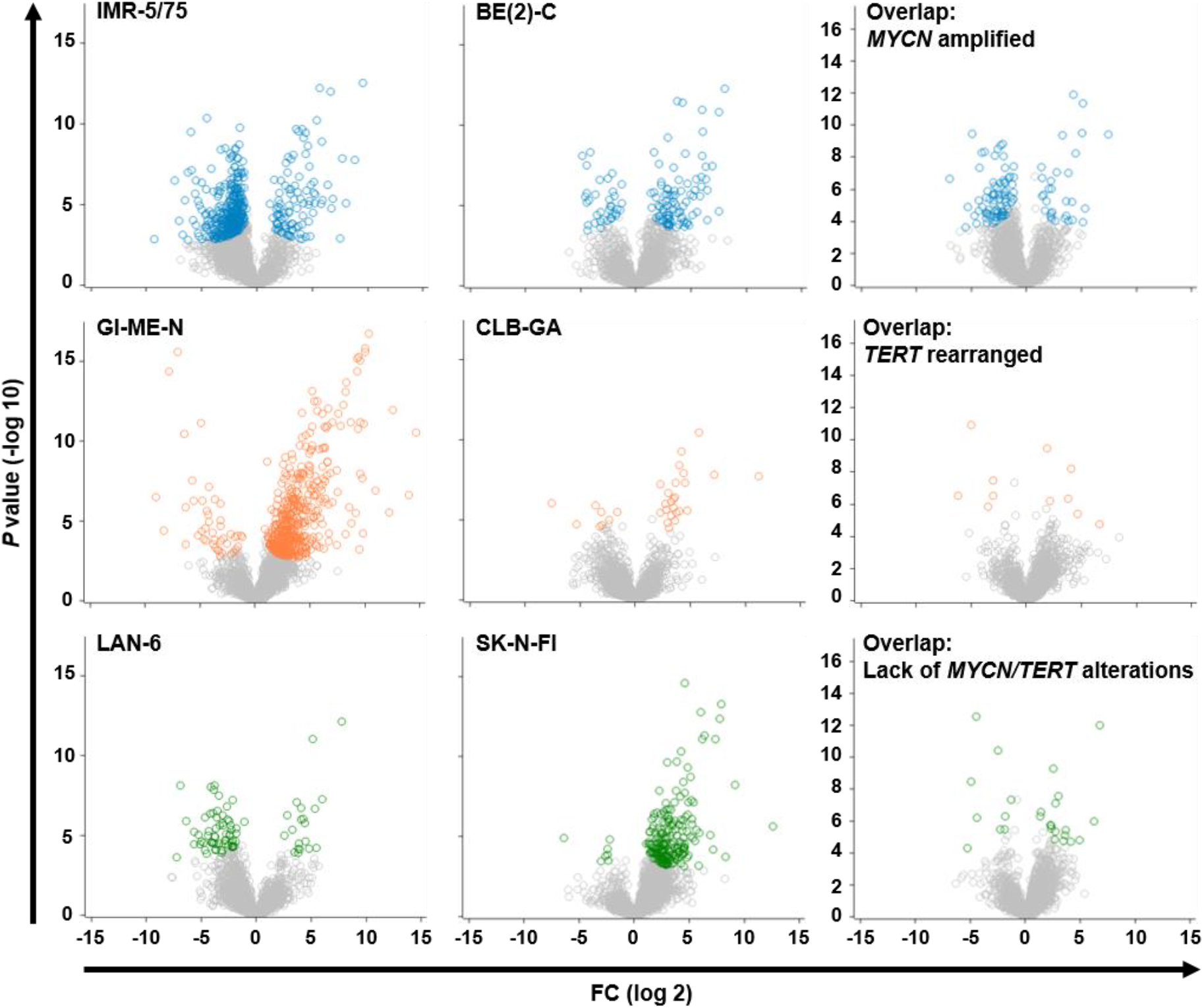
Genomic neuroblastoma subgroups share distinct sets of differentially abundant sEV proteins. Volcano plots showing differential protein abundance in sEVs derived from IMR-5/75 (477 DEPs), BE(2)-C (126 DEPs), GI-ME-N (521 DEPs), CLB-GA (31 DEPs), LAN-6 (85 DEPs) and SK-N-FI (213 DEPs) cells. Cell lines were analyzed in groups according to *MYCN* amplification (IMR-5/75 and BE(2)-C), *TERT* rearrangement (GI-ME-N and CLB-GA), or the absence of *MYCN* amplification and *TERT* rearrangement (LAN-6 and SK-N-FI), resulting in 110 DEPs (*MYCN* amplified), 11 DEPs (*TERT* rearranged) and 26 DEPs (lack of *MYCN*/*TERT* alternation). Colored circles indicate proteins with significant differential abundance (FDR < 1%) in the respective cell line, whereas grey circles indicate proteins not significantly different. Overlap plots show differential proteins in genomic subgroups.

To determine whether these subgroup-associated proteins exhibited coordinated abundance patterns across the complete sEV dataset, we next performed hierarchical clustering (**Fig. 3A**). Independent sEV preparations derived from the same neuroblastoma cell line showed highly similar protein-abundance profiles and clustered together, demonstrating the reproducibility of the EV proteomic signatures. Importantly, higher-order similarities were also apparent between cell lines belonging to the same genomic subgroup. sEVs derived from the *MYCN*-amplified IMR-5/75 and BE(2)-C cells displayed related protein-abundance patterns, whereas sEVs from the *TERT*-rearranged GI-ME-N and CLB-GA cells exhibited a distinct shared profile. LAN-6 and SK-N-FI, which lacked both *MYCN* amplification and *TERT* rearrangement, showed a third proteomic pattern. The hierarchical analysis further identified discrete modules of co-varying proteins with differential representation across the three genomic groups, as well as highly specific protein sets for each genomic group (**Fig. 3A**). Protein clusters showing relatively high abundance in one genomic group frequently exhibited lower abundance in the other groups, indicating coordinated differences in sEV cargo composition. Together with the overlap analysis in **Fig. 2**, these findings indicate that the relationship between sEV protein composition and genomic subgroup extends beyond individual differentially abundant proteins and involves broader patterns of protein abundance.

**Fig 3.**
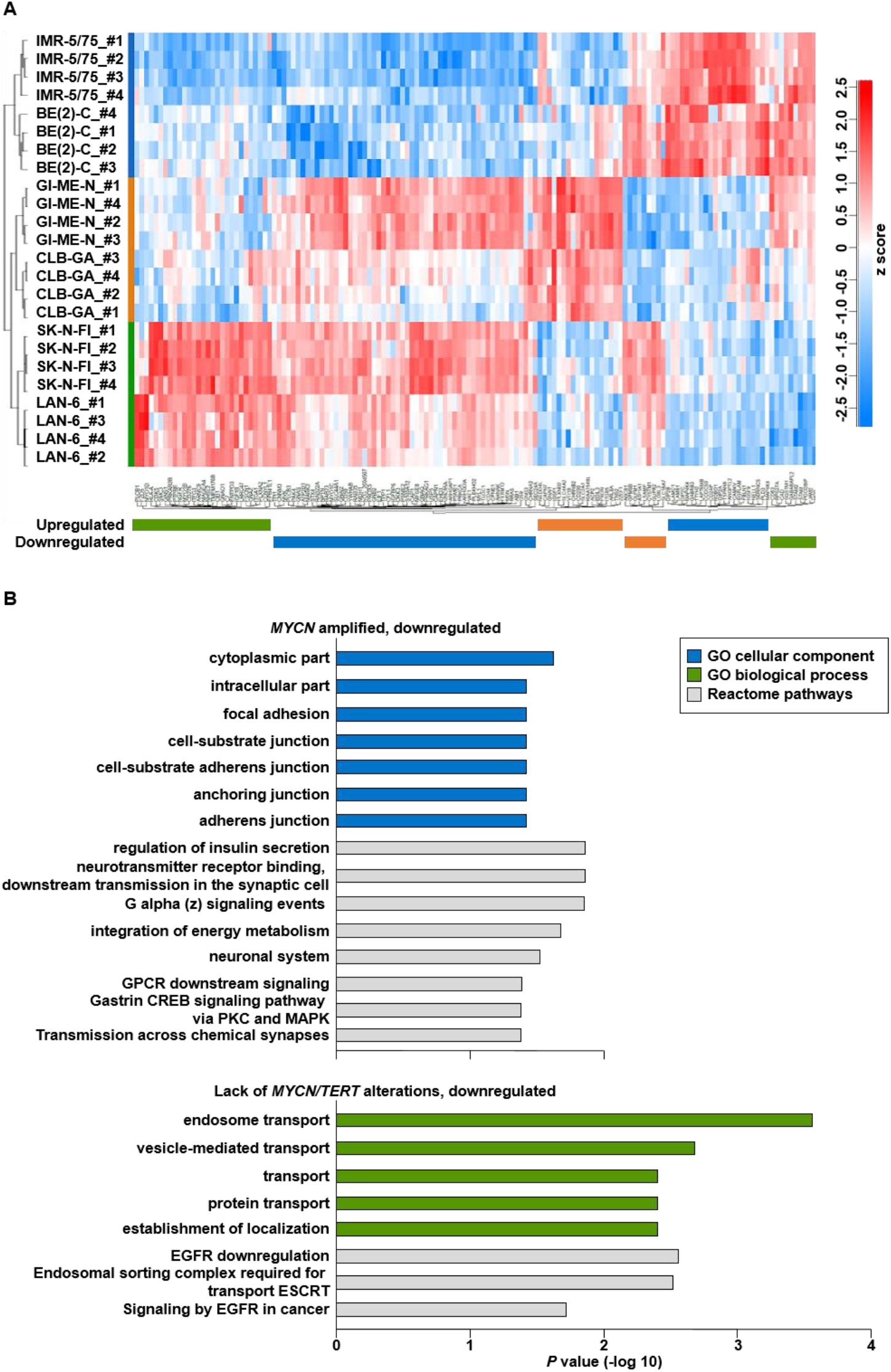
Subgroup-associated sEV protein signatures reveal coordinated proteomic patterns and functional pathways. **A**, Hierarchical clustering and heatmap of differentially abundant proteins (144 DEPs at FDR 1%) identified in sEVs derived from IMR-5/75, BE(2)-C, GI-ME-N, CLB-GA, LAN-6 and SK-N-FI cells. Rows represent individual sEV samples and columns represent proteins. Protein abundance is displayed according to the indicated color scale, with red and blue denoting higher and lower relative abundance, respectively. Dendrograms indicate hierarchical clustering of samples and proteins. Colored annotations indicate the genomic subgroup of the respective cell lines (*MYCN*-amplified, *TERT*-rearranged, or lacking *MYCN* amplification and *TERT* rearrangement). **B**, Functional enrichment analysis of proteins associated with the genomic subgroup-specific proteomic signatures. Bars show significantly enriched terms from Gene Ontology (GO) Cellular Component (blue), GO Biological Process (green) and Reactome pathway (grey) analyses. Bar length represents enrichment significance expressed as −log10(*P* value).

To explore the biological context of these subgroup-associated sEV protein signatures, functional enrichment analyses were performed using Gene Ontology (GO) and Reactome annotations (**Fig. 3B**). GO Cellular Component analysis identified significantly enriched cellular localizations among the subgroup-associated proteins, while GO Biological Process analysis identified biological processes represented within the respective protein signatures. Reactome analysis further identified pathways associated with these protein sets. The enrichment of multiple functional categories and pathways indicates that the proteomic differences between the genomic groups involve functionally related protein networks rather than isolated changes in individual sEV proteins. Collectively, the differential abundance, hierarchical clustering and functional enrichment analyses demonstrate that the sEV proteome contains reproducible signatures reflecting the genomic characteristics of the parental neuroblastoma cells. sEVs derived from *MYCN*-amplified, *TERT*-rearranged and *MYCN*/*TERT*-unaltered neuroblastoma models exhibited distinct patterns of protein abundance and functional enrichment. These findings indicate that the molecular heterogeneity defining neuroblastoma subgroups is also reflected in the protein composition of tumor cell-derived sEVs.

### Parental cell state is reflected in the proteomic composition of neuroblastoma-derived sEVs

Having identified genomic subgroup-associated signatures within the sEV proteome (**Fig. 2** and **Fig. 3**), we next investigated how sEV protein composition relates to the proteomic state of the corresponding parental neuroblastoma cells. To this end, we analyzed the proteomes of the six neuroblastoma cell lines, covering 2850 proteins quantified in both, the parental cell and their respective sEVs (**Fig. 4**). Enrichment values (log2 fold changes from the “one against all others” comparisons) for both, sEVs and parental cells, were used for joined principal component analysis (PCA). Data distribution revealed a marked distinction between the mesenchymal and adrenergic neuroblastoma models (**Fig. 4A**). GI-ME-N, the mesenchymal cell line within the proteomic discovery panel, separated from the adrenergic cell lines. Notably, this distinction was apparent for both the parental GI-ME-N cells and their corresponding sEVs, whereas the parental cells and sEVs derived from the adrenergic models occupied distinct regions from GI-ME-N in the principal component space. PC1 and PC2 accounted for 24.9% and 15.3% of the total variance, respectively. These findings suggested that proteomic features associated with the mesenchymal state of GI-ME-N cells are also represented within their sEV protein cargo.

**Fig 4.**
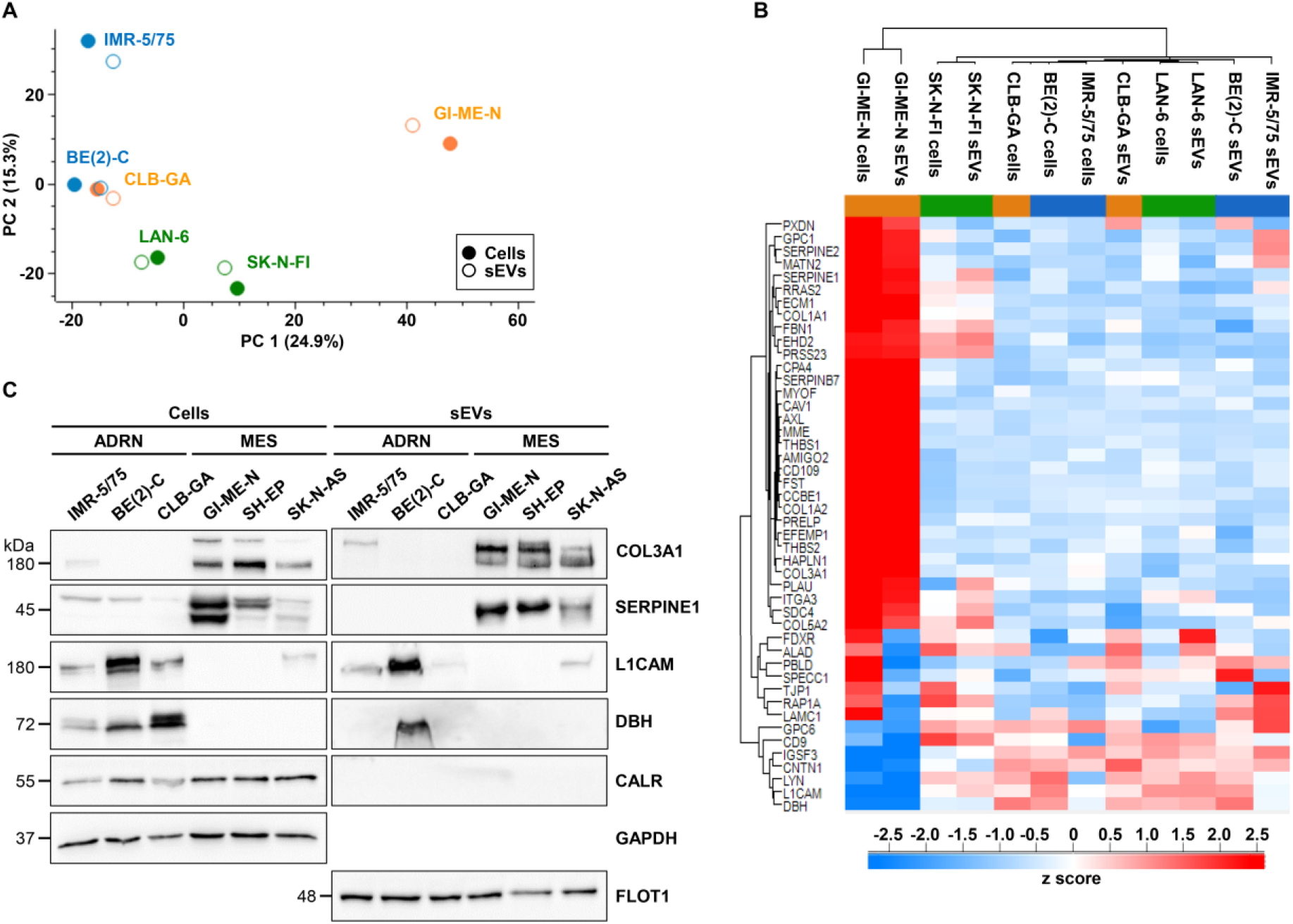
Mesenchymal and adrenergic neuroblastoma cell states are reflected in parental cell and sEV proteomes. **A**, Principal component analysis of proteomic profiles from neuroblastoma parental cells and their sEVs. Filled and open symbols indicate parental cells and sEVs, respectively. Colours denote the respective neuroblastoma cell lines. PC1 and PC2 explain 24.9% and 15.3% of the total variance, respectively. **B**, Hierarchical clustering and heatmap of proteins, significantly different at FDR < 5% in parental cells and corresponding sEVs from IMR-5/75, BE(2)-C, CLB-GA, GI-ME-N, LAN-6 and SK-N-FI. Protein enrichment is displayed as *z*-scores according to the indicated colour scale. Rows indicate proteins and columns indicate parental cell or corresponding sEV samples. **C**, Immunoblot analysis of adrenergic (ADRN)- and mesenchymal (MES)-associated proteins in parental cells and corresponding sEVs from IMR-5/75, BE(2)-C, CLB-GA and GI-ME-N, together with the additional mesenchymal neuroblastoma cell lines SH-EP and SK-N-AS. L1CAM anddopamine β-hydroxylase (DBH) were analysed as ADRN-associated proteins, and SERPINE1 and COL3A1 as MES-associated proteins. CALR, GAPDH and FLOT1 were analysed as indicated. Molecular masses are shown in kDa.

Hierarchical clustering of significantly different proteins (FDR < 5%) in sEVs and parental cells further supported the relationship between parental cell state and sEV protein composition (**Fig. 4B**). GI-ME-N cells and GI-ME-N-derived sEVs clustered separately from the corresponding cell and EV samples of the adrenergic neuroblastoma models. The separation was associated with coordinated differences across a broad set of proteins, including proteins linked to adrenergic and mesenchymal neuroblastoma cell states. In particular, GI-ME-N cells and their sEVs exhibited a distinct abundance pattern compared with the adrenergic models, showing that the phenotypic differences between the parental cells can be reflected in the sEV cargo composition.

Because GI-ME-N represented the only mesenchymal cell line in the proteomic discovery cohort, we next investigated whether the observed protein pattern was restricted to GI-ME-N or was also present in additional mesenchymal neuroblastoma models. We therefore extended the analysis to the mesenchymal cell lines SH-EP and SK-N-AS and examined selected adrenergic- and mesenchymal-associated proteins by immunoblotting in parental cells and their corresponding sEVs (**Fig. 4C**). The expanded panel included the adrenergic models IMR-5/75, BE(2)-C and CLB-GA and the mesenchymal models GI-ME-N, SH-EP and SK-N-AS.

Immunoblot analysis of the adrenergic-associated proteins L1CAM and dopamine β-hydroxylase (DBH) and the mesenchymal-associated proteins SERPINE1 and COL3A1 revealed cell-state-associated protein patterns across the expanded panel. Importantly, SH-EP and SK-N-AS showed patterns similar to those observed for GI-ME-N, providing independent support for the distinction initially identified in the proteomic analysis. Corresponding sEV preparations displayed related differences in these markers, indicating that protein features associated with the adrenergic or mesenchymal state of the parental cells are also represented in their sEV cargo. CALR, GAPDH and the sEV-associated protein FLOT1 were additionally assessed in the respective cell and sEV preparations.

Collectively, these findings reveal an additional level of organization of the neuroblastoma sEV proteome. While analysis of sEV cargo alone identified signatures associated with *MYCN* amplification and *TERT* rearrangement (**Fig. 2, Fig. 3**), integration of the sEV and parental-cell proteomes revealed an additional association with the adrenergic–mesenchymal cell state. The concordant protein patterns observed in the additional mesenchymal models SH-EP and SK-N-AS further support this relationship beyond GI-ME-N. Thus, neuroblastoma sEVs retain proteomic features associated with the adrenergic or mesenchymal state of their parental cells, indicating that EV cargo reflects both genomic and phenotypic heterogeneity of the tumor cells from which they originate.

## DISCUSSION

Our high-resolution proteomic profiling of sEVs derived from neuroblastoma cell lines represents a comprehensive, side-by-side analysis of paired cell-sEV proteomes across distinct genetic and phenotypic neuroblastoma subtypes. A central finding of this study is that neuroblastoma-derived sEVs carry robust, highly reproducible protein signatures that are systematically coordinated with both the genomic landscape and the differentiation state of the parent cell. This suggests that the sEV release and packaging machineries are not stochastic, but are instead governed by the core genetic programs^9,11,12^ and epigenetic circuitries^14,16^ that dictate neuroblastoma pathogenesis and lineage identity.

The distribution of the sEV proteomes into distinct clusters mirrors the clinically decisive genomic aberrations of high-risk neuroblastoma, specifically *MYCN* amplification and *TERT* rearrangement. The sEVs from *MYCN*-amplified cell lines (IMR-5/75 and BE(2)-C) and *TERT*-rearranged lines (GI-ME-N and CLB-GA) exhibit distinctive, overlapping protein profiles within their respective subgroups, suggesting that subgroup-specific transcriptomic programs govern the cargo of secreted vesicles. This is consistent with the model of high-risk neuroblastoma, where transformation arises at specific sympathoadrenal progenitor stages held open by core transcriptional networks.^37^ The coordinated sorting of oncoproteins and metabolic regulators into sEVs underscores their role as functional extensions of the parental phenotype.^4^

At the level of phenotypic plasticity, we observed a concordance between the ADRN versus MES transdifferentiation states of the parental cells and their corresponding sEV cargo. The presence of classic ADRN markers such as L1CAM and DBH in ADRN sEVs stands in sharp mutually exclusive contrast to the enrichment of MES-associated markers, SERPINE1 and type III collagen (COL3A1) in sEVs from MES cell models. Cellular plasticity and lineage transdifferentiation represent critical non-genetic mechanisms of therapeutic resistance and disease progression in patients at high-risk. Prior studies have established that MES cells exhibit enhanced migratory capacity, stemness, and resistance to standard chemotherapy.^18^ Our demonstration that these functional phenotypic states are preserved and readable within the sEV proteome highlights their potential as non-invasive liquid biopsy biomarkers to track plastic lineage trajectories in real-time.^38^ This aligns with a recent study by Lampis and colleagues^17^, which identified microRNA signatures such as miR-199a-3p and let-7f-5p in the MES phenotype both in patient tissue and in sEVs to upregulate key mesenchymal players such as FN1, CD44, and YAP1.

Neuroblastoma-derived sEVs mediate premetastatic niche formation in preferred secondary organs, thus acting as active, long-range intercellular messengers.^4,39^ The deposition of MES-associated sEV proteins such as SERPINE1 and COL3A1 contributes to extracellular matrix remodeling, stromal cell activation and vascular permeability in distant tissues, preparing a supportive soil for disseminating tumor cells.^4^ Wills and colleagues recently showed that doxorubicin-treated neuroblastoma cells secrete sEVs enriched in PTX3 and PLAT that accelerate hepatic pre-metastatic niche priming by inducing fibronectin accumulation and CD45+ myeloid cell infiltration.^5^ Fietta and colleagues utilized a transgenic zebrafish xenotransplantation model to track the in vivo fate of neuroblastoma sEVs, demonstrating that hypoxic sEVs are rapidly internalized by endothelial cells, stimulate angiogenesis in the sub-intestinal veins, and mobilize host macrophages to home in the caudal hematopoietic tissue, the zebrafish functional analog of the mammalian bone marrow.^6^ At the molecular level, the hypoxic sEV-mediated pre-conditioning triggers a significant upregulation of matrix metalloproteinase-9 and the inflammatory chemokine cxcl8b in the hematopoietic niche, culminating in accelerated tumor cell proliferation and metastatic outgrowth.^6^ Further, matrix- and protease-associated sEV cargo has been reported to act on the endothelium, targeting tight-junction proteins such as ZO-1 and claudin-5 to increase vascular permeability^40^, promote angiogenesis^41^ and drive endothelial branching and perivascular inflammation^42^.

Neuroblastoma-derived sEVs also exert local and systemic immunomodulatory effects to shield colonizing tumor cells from host surveillance.^4^ Our results indicate that sEVs may participate in a complex immunoregulatory network within the metastatic bone marrow microenvironment. Marimpietri and colleagues showed that sEVs isolated directly from the bone marrow of neuroblastoma patients exhibit a marked upregulation of immune checkpoint molecules such as HLA-G and PD-L1, which actively suppress CD4+ and CD8+ T-cell proliferation.^7^ Furthermore, these bone marrow sEVs modulate local inflammatory states by suppressing the secretion of interleukin-6 and interferon-α from activated mononuclear cells while simultaneously enhancing the release of GM-CSF.^7^ Within the bone marrow niche, sEVs also interact dynamically with resident mesenchymal stromal cells to induce pro-tumorigenic cytokine secretion (such as IL-6, IL-8, and VEGF) via AKT/ERK phosphorylation and promote osteogenic transdifferentiation,^43,44^ thereby transforming the bone marrow into an immunologically privileged, tumor-supportive environment.

The nature of sEVs to harbor and transport genetic, transcriptomic and proteomic features of their parent cells may provide novel avenues for precision oncology in pediatric patients. The isolation of circulating sEVs from longitudinal plasma samples may evolve as a powerful, minimally invasive tool to monitor real-time tumor dynamics and detect therapy-induced lineage transdifferentiation.^18,38^ Moreover, targeting sEV-mediated signaling networks may provide novel therapeutic opportunities. Preclinical studies have shown that blocking sEV biogenesis or secretion using farnesyltransferase inhibitors (e.g., tipifarnib) or neutral sphingomyelinase inhibitors (e.g., GW4869) rescues host natural killer cell recruitment and synergistically enhances the therapeutic efficacy of standard chemotherapies and anti-GD2 immunotherapy in neuroblastoma models.^4^

Several considerations should be kept in mind, which outline clear pathways for future investigation. First, while utilizing two representative cell lines per genomic subgroup enabled an informative profiling of shared traits, subgroup-specific properties and individual cell line identities remain closely linked. To fully decouple these factors and confirm that the observed overlaps are driven primarily by *MYCN* amplification or *TERT* rearrangement, validation in larger panels represents an essential next step. Second, within the proteomic discovery panel, the mesenchymal state is represented by a single model (GI-ME-N). The highly concordant marker patterns observed in independent models (SH-EP and SK-N-AS) provide strong, external validation for our cell-state interpretation. Third, sEVs were harvested under standardized, normoxic and drug-free culture conditions using EV-depleted serum to establish a clean and reproducible molecular baseline. While this design excludes the acute microenvironmental or therapeutic pressures seen in treated patients, it provides a crucial reference dataset from which future studies can measure translational and treatment-induced changes. Finally, this study was designed to comprehensively characterize the molecular cargo of neuroblastoma sEVs, rather than their downstream functional consequences. While recipient cell uptake and functional assays were outside the scope of this work, our cell-state-resolved proteomic profiles deliver a highly curated candidate list that serves as the necessary foundation for future functional experiments in endothelial and stromal microenvironments.

In summary, this study analyzed the sEV cargo using proteomic approaches, which demonstrated that sEVs cluster according to their donor cell line and reflect underlying tumor cell states. Notably, sEV proteomes allowed discrimination between adrenergic (ADRN) and mesenchymal (MES) neuroblastoma states, with MES-associated sEVs enriched for proteins linked to extracellular matrix organization and tissue remodeling (e.g. COL3A1, SERPINE1), suggesting a potential role in vascular niche modulation.

## ACKNOWLEDGEMENTS

The authors thank Jasmin Wünschel for excellent technical support and Nicole Huebener and Rogier Versteeg (Department of Pediatric Oncology and Hematology, Charité – Universitätsmedizin Berlin) for helpful discussions and critical feedback on the manuscript. The authors acknowledge the use of Gemini Notebook (Google) solely for assistance in shaping and editing the manuscript text, while the final intellectual content and scientific accuracy remain the sole responsibility of the authors.

