## Supplementary Data file for "Neuroblastoma-derived small extracellular vesicles retain cellular identity and adrenergic/mesenchymal state signatures"

**Table of Contents**

**Supplementary Figures** starting p. 2

**Legends to Supplementary Figures** starting p. 4

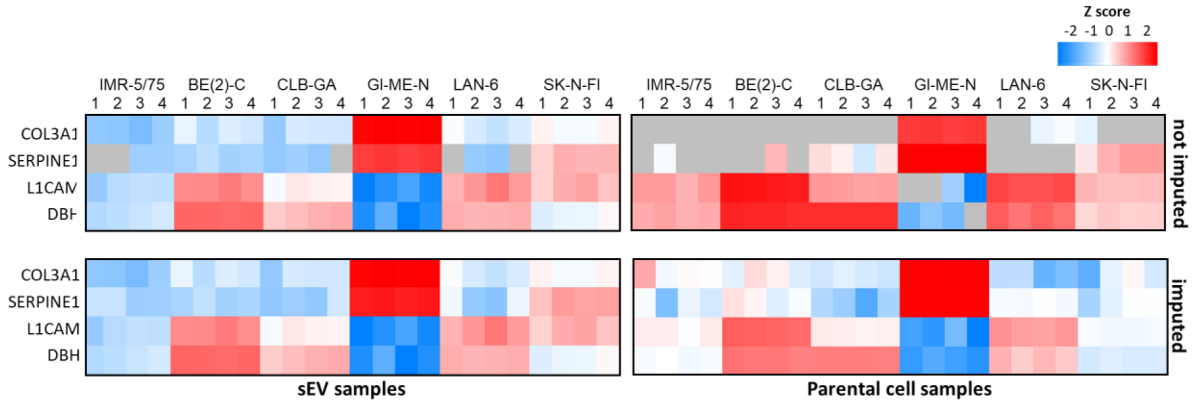


**Suppl. Fig. S1. Imputation preserves relative abundance patterns of mesenchymal markers in paired parental cell and sEV samples.** Heatmaps of abundance Z-scores of mesenchymal marker proteins in sEVs (*left panels*) and parental cells (*right panels*). Shown are non-imputed (*top panels*) and imputed data (*bottom panels*) for all cell lines and the respective paired sEV samples. Colors represent scaled expression values. Blue, low expression values; red, high expression levels; grey, not detected (below detection limit).

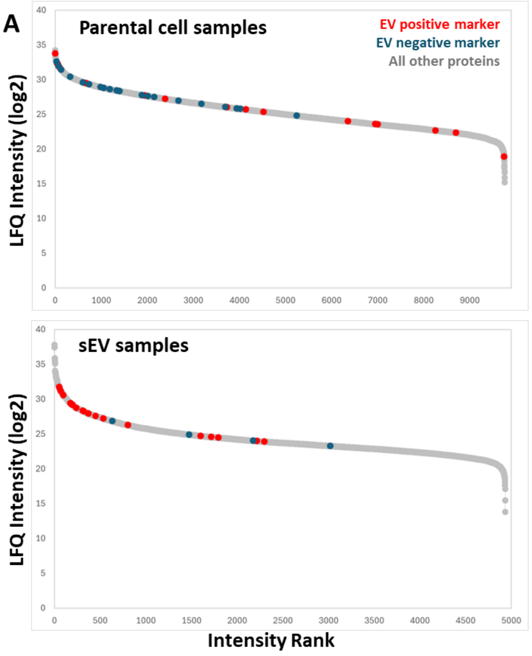


**B**

**sEV positive marker**

Intensity (log2) 20 25 30 35

| Gene names | Median log2 LFQ intensity sEVs | Median log2 LFQ intensity parental cells |
|---|---|---|
| CD63 | 28.72 | 26.08 |
| CD81 | 28.26 | 25.70 |
| CD9 | 30.58 | 26.04 |
| ENO1 | 29.20 | 31.96 |
| FLOT1 | 28.33 | 27.72 |
| GAPDH | 29.42 | 32.24 |
| HSP90AA1 | 27.61 | 31.75 |
| PDCD6IP | 31.63 | 29.47 |
| PKM | 28.74 | 29.50 |
| SDCBP | 31.75 | 25.86 |
| TSG101 | 29.41 | 27.26 |
| TSPAN14 | 27.92 | 22.70 |
| TSPAN15 | 24.48 | 23.60 |
| TSPAN18 | 26.29 | 24.03 |
| TSPAN3 | 27.23 | 25.38 |
| TSPAN33 | 23.91 | n.d |
| TSPAN9 | 24.73 | 18.91 |
| YWHAZ | 29.26 | 32.35 |

**sEV negative marker**

| Gene names | Median log2 LFQ intensity sEVs | Median log2 LFQ intensity parental cells |
|---|---|---|
| AGO1 | n.d | 25.79 |
| BCL2 | n.d | 25.86 |
| CALR | 24.03 | 31.47 |
| CANX | 24.92 | 31.99 |
| COA3 | n.d | 27.78 |
| COA7 | n.d | 26.96 |
| COX17 | n.d | 26.07 |
| COX19 | n.d | 24.82 |
| COX20 | n.d | 26.52 |
| COX4I1 | n.d | 29.63 |
| COX5A | n.d | 27.64 |
| COX5B | n.d | 29.35 |
| COX6B1 | n.d | 28.82 |
| COX6C | n.d | 28.44 |
| CYCS | 23.30 | 28.82 |
| GOLGA2 | n.d | 28.39 |
| HSP90B1 | 26.86 | 32.63 |
| MT-CO2 | n.d | 30.41 |
| TOMM20 | n.d | 27.50 |
| TOMM22 | n.d | 28.93 |
| TOMM40 | n.d | 28.36 |
| TOMM7 | n.d | 28.63 |

**Suppl. Fig. S2. Proteomic evaluation of sEV purity. A**. Protein intensity rank (x-axis) based on the median log2 LFQ intensity values (y-axis) for all parental cells (top) and sEVs (bottom). Proteins of selected purity marker groups are color-coded (red, sEV positive marker; blue, sEV negative marker; grey, all other proteins). **B**. Overview of intensity values of positive (*left*) and negative (*right*) sEV marker proteins, color-scaled to the range of observed LFQ intensities within either sEVs or paternal cells. n.d, protein was not detected.

### LEGENDS TO SUPPLEMENTARY FIGURES

**Suppl. Fig. S1. Imputation preserves relative abundance patterns of mesenchymal markers in paired parental cell and sEV samples.** Heatmaps of abundance Z-scores of mesenchymal marker proteins in sEVs (*left panels*) and parental cells (*right panels*). Shown are non-imputed (*top panels*) and imputed data (*bottom panels*) for all cell lines and the respective paired sEV samples. Colors represent scaled expression values. Blue, low expression values; red, high expression levels; grey, not detected (below detection limit).

**Suppl. Fig. S2. Proteomic evaluation of sEV purity. A**, Protein intensity rank (x-axis) based on the median log2 LFQ intensity values (y-axis) for all parental cells (top) and sEVs (bottom). Proteins of selected purity marker groups are color-coded (red, sEV positive marker; blue, sEV negative marker; grey, all other proteins). **B**, Overview of intensity values of positive (*left*) and negative (*right*) sEV marker proteins, color-scaled to the range of observed LFQ intensities within either sEVs or paternal cells. n.d, protein was not detected.
